# The Comet Toolbox: Improving Robustness in Network Neuroscience Through Multiverse Analysis

**DOI:** 10.1101/2024.01.21.576546

**Authors:** Micha Burkhardt, Carsten Gießing

## Abstract

In network neuroscience, a broad range of methods for estimating dynamic functional connectivity from fMRI data and subsequent analyses using graph-theoretic approaches have been introduced in recent years. However, in the absence of ground truths regarding the validity of analytical steps in capturing true brain dynamics, researchers are often faced with a multitude of arbitrary, yet defensible, analytical choices, raising concerns regarding the robustness of results. Here, we aim to address this issue by implementing a comprehensive suite of dynamic functional connectivity methods in a unified Python software package, allowing for a diverse exploration of brain dynamics. Anchored in the framework of multiverse analysis, the present work introduces a workflow for systematically exploring different methodological choices. The developed toolbox includes a graphical user interface for ease of use and accessibility for those who wish to operate outside of a script-based pipeline. Comprehensive documentation and demo scripts are included to support adoption and usability. By promoting transparency and robustness, Comet aims to promote best practice in the study of brain dynamics.

## 1 Introduction

In recent years, the field of network neuroscience has witnessed a rapid expansion in methodological diversity, particularly in the estimation of dynamic functional connectivity (dFC) and its application to graph-theoretic analyses. While this growing variety of approaches reflects methodological progress, it has also introduced significant uncertainty regarding the robustness and reproducibility of findings. This issue is not unique to neuroscience: across fields such as psychology, economics, and genomics, it has become evident that arbitrary or under-specified analysis choices can critically shape results and limit replicability (Frias-Navarro et al., 2020; Ioannidis, 2005; Simmons et al., 2011). Despite increasing efforts to promote transparency, for example through open data, preregistration, or more rigorous reporting standards, a general replication crisis persists in the scientific literature (Fanelli, 2018; Frias-Navarro et al., 2020). Within neuroimaging, these concerns are amplified by the inherently noisy nature of the data and the large number of decisions required at each step of analysis. As highlighted by recent large-scale collaborative studies, different analysis teams applying different pipelines to the same dataset can arrive at divergent conclusions (Botvinik-Nezer et al., 2020), underscoring the importance of methodological transparency and robustness.

Dynamic functional connectivity aims to characterise time-varying patterns of functional interactions between brain regions, offering a window into the brain’s dynamic reconfiguration across cognitive states and in response to internal or external stimuli (Hutchison et al., 2013; Iraji et al., 2021). It has shown promise in elucidating mechanisms underlying a range of neuropsychiatric and neurological disorders, including schizophrenia, Alzheimer’s disease, and mood disorders (Calhoun et al., 2014; Gonzalez-Castillo & Bandettini, 2018). Numerous approaches have been proposed to estimate dFC, such as sliding-window correlation, co-activation patterns, time-varying graphical models, and Hidden Markov Models (Hutchison et al., 2013; Lurie et al., 2020; Preti et al., 2017). Yet, despite these advances, substantial variability remains across methods, and no consensus has emerged on best practices for estimating dFC (Thompson et al., 2018; Torabi et al., 2024). Moreover, test-retest studies have reported low-to-moderate reliability of dFC estimates, with intraclass correlation coefficients often falling between 0.2 and 0.5, raising important concerns about their stability and interpretability (Choe et al., 2017; Zhang et al., 2018). These findings highlight the need for systematic tools that can assess the impact of analytic flexibility on conclusions drawn from dynamic functional connectivity analyses.

To address these challenges, we introduce Comet - a user-friendly and open-source Python toolbox for multiverse analysis designed specifically for neuroimaging research (Figure 1). As a recently proposed approach to improve robustness and transparency in data analysis, multiverse analysis involves the systematic evaluation and reporting of all defensible combinations of analytic decisions, from data pre-processing to statistical modelling, rather than relying on a single, potentially arbitrary pipeline (Steegen et al., 2016). This approach provides a principled framework to quantify the influence of methodological decisions on observed outcomes and enables a transparent overview of the robustness of reported effects (Dafflon et al., 2022). In this context, Comet offers several key advantages over existing packages. It supports a wide range of methods for estimating dynamic functional connectivity, including both established and novel approaches, and allows users to construct and compare entire analysis pipelines. Beyond dFC, Comet’s modular architecture permits the incorporation of user-defined methods, making it adaptable to a variety of neuroimaging modalities and research questions. Designed for both novice and experienced users, the toolbox is accessible via a graphical user interface (GUI) for ease of use, as well as through Python scripting for integration into reproducible, automated workflows. Comet further includes extensive learning resources, such as example pipelines, tutorials, and visual reporting tools, to guide users through each stage of the multiverse process. These features make the toolbox particularly valuable not only for methodological research but also for education and reproducible reporting in applied neuroscience. Development of Comet is ongoing, ensuring that the toolbox will continue to evolve with new features and methodological updates to further support transparent, flexible, and rigorous neuroimaging analyses.

**Figure 1:**
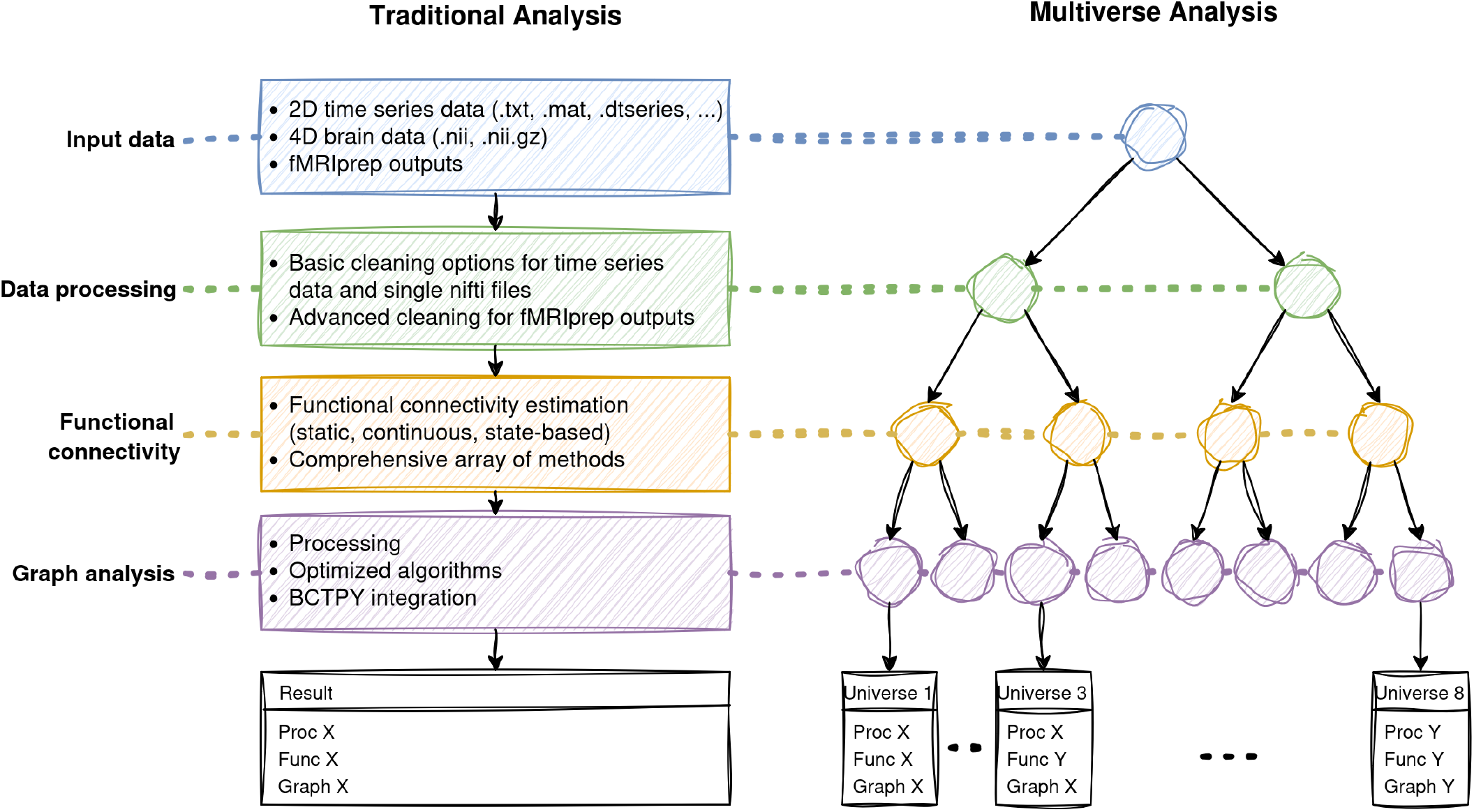
Features of the Comet Toolbox. Comet incorporates a broad set of functionalities spanning input handling, data processing, (dynamic) functional connectivity estimation, and graph-theoretic analysis. On the left, a traditional single-pipeline analysis is illustrated, where a single result is produced from a fixed sequence of choices, with available features of the toolbox listed in each box. On the right, the multiverse analysis framework is shown, illustrating possible combinations of defensible analytic decisions across the pipeline. Each branching point represents a user-specified set of alternatives at a particular analysis stage. The toolbox supports multiple input formats and includes data cleaning and preprocessing options. Connectivity estimation covers static, continuous, and state-based approaches, with a broad range of algorithms available. The graph module includes tools for network processing (e.g., thresholding, binarisation), popular graph-theoretic measures, and integration with the Python implementation of the Brain Connectivity Toolbox. In the Result boxes, each entry reflects a unique combination of processing, connectivity, and graph analysis choices, referred to as a universe in the multiverse framework. Users can also integrate custom methods at any step, ensuring full extensibility.

The software is publicly available at: https://github.com/mibur1/comet. In the sections that follow, we detail the architecture and core functionalities of Comet, provide illustrative applications to multi-verse analysis in dFC, and outline future directions aimed at broadening its utility for the neuroimaging community.

## 2 Methods

The main contribution of this work is an open source Python toolbox for multiverse analysis, with a focus on network neuroscience. The toolbox provides both a graphical user interface (Figure 2) and a flexible scripting API (Figure 5, 6) to accommodate diverse workflows. A brief overview of the included features and methods will be provided in the following subsections.

**Figure 2:**
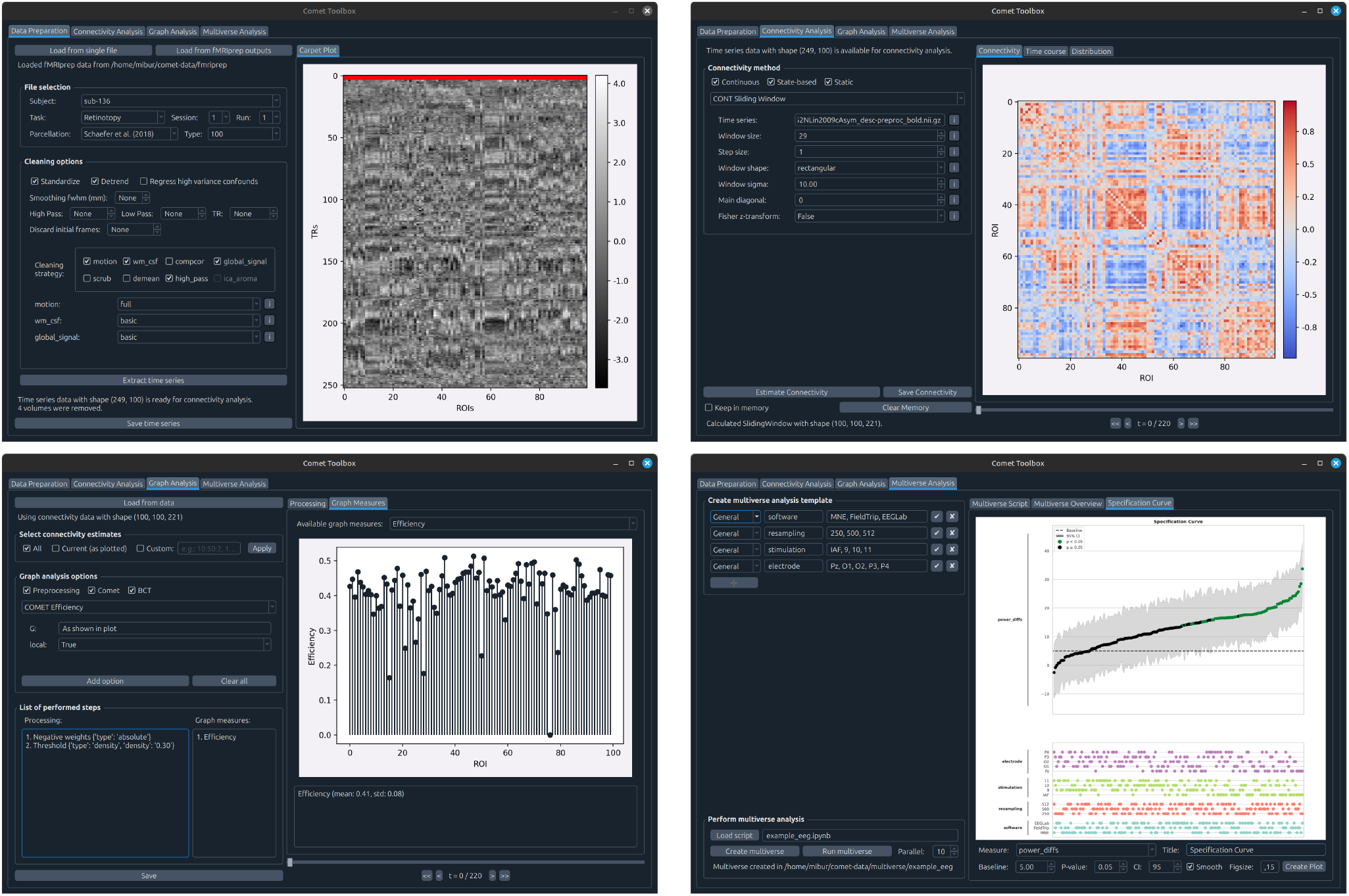
Graphical user interface. Comet offers a graphical user interface for most of its features, including data handling and processing (top left), functional connectivity estimation (top right), graph analysis (bottom left), and multiverse analysis (bottom right). All features can be used individually or in combination, allowing for a flexible usage.

### 2.1 Functional Connectivity

The estimation of functional connectivity can be broadly divided into three distinct classes of methods, based on how temporal variations are addressed: Static functional connectivity, continuously varying dynamic functional connectivity, and state-based dynamic functional connectivity.

*Static functional connectivity* methods assume that connectivity remains constant throughout the duration of the recording, yielding a single overall connectivity estimate (Van Den Heuvel & Pol, 2010). In contrast, dynamic approaches aim to capture temporal fluctuations in connectivity (Iraji et al., 2021).

*Continuously varying dynamic functional connectivity* methods estimate connectivity at each time point (or a specified set of time points) and can further be divided into two sub-classes. Amplitude-based methods operate directly in the time domain of the signal, deriving connectivity estimates from the amplitude of BOLD time series data. Alternatively, time-frequency-based methods incorporate both frequency and phase information, revealing insights into temporal synchronization and rhythmic coordination between brain regions (see, for example, Lurie et al. (2020)).

*State-based dynamic functional connectivity* methods characterise brain connectivity as a sequence of transitions between a finite set of discrete states, each assumed to exhibit quasi-stationary connectivity patterns. Transitions between these states reflect dynamic, time-dependent reconfigurations of the underlying functional network.

Comet currently includes 18 different methods spanning these categories as summarized in Table 1. The methodological diversity exists not only between different approaches but also within individual methods, for example, in the choice of hyper-parameters such as the sliding window size or the number of connectivity states making them an ideal subject for multiverse analysis.

**Table 1:** Functional connectivity methods included in the Comet toolbox.^a^

| Type | Method | Domain | Reference |
| --- | --- | --- | --- |
| Static | Pearson Correlation | Amplitude | – |
|  | Partial Correlation | Amplitude | – |
|  | Mutual Information | Amplitude | – |
| Continuous | (Tapered) Sliding Window Correlation | Amplitude | – |
|  | Flexible Least Squares | Amplitude | Liao et al. (2014) |
|  | Multiplication of Temporal Derivatives | Amplitude | Shine et al. (2015) |
|  | Spatial Distance | Amplitude | Thompson et al. (2017) |
|  | Jackknife Correlation | Amplitude | Richter et al. (2015) |
|  | Edge-centric Connectivity | Amplitude | Faskowitz et al. (2020) |
|  | Dynamic Conditional Correlation | Amplitude | Lindquist et al. (2014) |
|  | Phase Synchronization | Time-freq | Honari et al. (2021) |
|  | Wavelet Coherence | Time-freq | Billings et al. (2021) |
|  | Leading Eigenvector Dynamics | Time-freq | Cabral et al. (2017) |
| State-based | Sliding Window Clustering | Amplitude | Allen et al. (2014) |
|  | Co-activation Patterns | Amplitude | X. Liu and Duyn (2013) |
|  | Discrete Hidden Markov Model | Amplitude | Vidaurre et al. (2017) |
|  | Continuous Hidden Markov Model | Amplitude | Ou et al. (2015) |
|  | Windowless | Amplitude | Yaesoubi et al. (2018) |

<sup>a</sup> For a full list of parameters and examples, refer to the toolbox documentation.

### 2.2 Graph Analysis

Comet offers a range of graph-analysis techniques for estimating network topology within a multiverse analysis framework. To facilitate this, it provides wrapper functions for most of the algorithms available in the Python implementation of the Brain Connectivity Toolbox (Rubinov & Sporns, 2010). Further, for performance related reasons, Comet also offers custom implementations for computationally heavy algorithms such as local efficiency, which compile to machine code at run time using the Numba library (Lam et al., 2015). This significantly increases calculation speed, which is particularly helpful when dealing with large graphs. An example for this is shown for the calculation of local efficiency in a relatively large network containing 379 nodes, in which performance is increased by a factor of 3 to 10 compared to previous implementations of the method (Figure 3). A complete list of the currently included methods can be found in the documentation of the toolbox.

**Figure 3:**
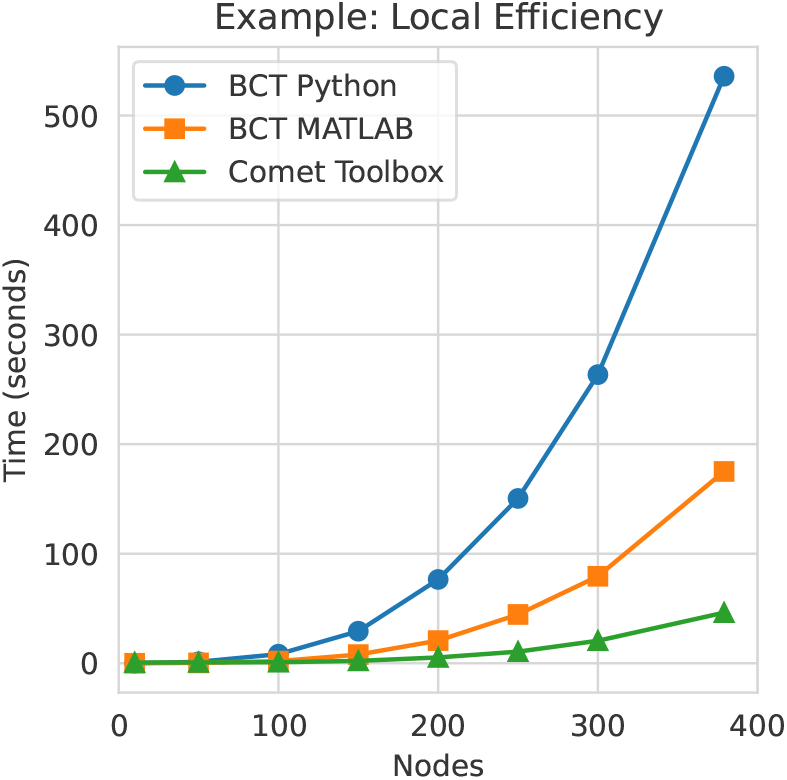
Toolbox performance. Implementations of computationally heavy graph measures included in Comet compile to native machine code at run time, which significantly improves performance. For example, on a laptop equipped with an AMD Ryzen 7 PRO 7840U, the calculation of local efficiency for a weighted brain network with 379 nodes at 50% density shows improved calculation speed compared to the Python version (*>*10x) and MATLAB version (*>*3x) of the Brain Connectivity Toolbox (BCT).

### 2.3 Multiverse Analysis

Comet implements a flexible and modular framework for conducting multiverse analyses. Users specify the possible alternatives for each analytic decision, thereby defining the forking paths within their analysis pipeline. The toolbox then automatically generates analysis scripts for all possible combinations of these decision options (or for a selected subset based on pruning rules). Each unique combination of decisions constitutes a separate universe within the multiverse framework, representing one defensible analytic pathway from data processing and analysis to result. Universes can then be evaluated and their outcomes are aggregated to assess the robustness of findings. This design allows users to easily inspect individual scripts and provides the flexibility to execute them either sequentially or in parallel, depending on the available computational resources. A brief illustration of the multiverse workflow is provided in Figure 6 and will be discussed in more detail in Section 3.2.

## 3 Usage

The Comet toolbox can be installed via the Python Package Index (PyPI) using pip. As this will automatically install or update required packages, we strongly recommend users to set up a dedicated environment (e.g., venv^1^ or conda^2^) to avoid potential version conflicts (4).

A key design goal of the toolbox is to support flexible and modular use. Researchers do not need to run their entire analysis pipeline within Comet. Instead, the toolbox can be used as a standalone tool to estimate dynamic functional connectivity or to conduct multiverse analyses. This makes it suitable for a wide range of study designs and research questions. At the time of this publication, Comet includes five main modules, which are described in detail in the documentation:

`comet.connectivity` (Dynamic) functional connectivity methods,

`comet.graph` Graph analysis functions,

`comet.multiverse` Multiverse analysis functions,

`comet.utils` Data loading and processing utilities,

`comet.cifti` CIFTI parcellation and file handling.

This modular design allows users to integrate individual components of the Comet toolbox, such as specific functional connectivity estimation methods, into their own analysis pipelines with minimal changes. For instance, users can apply Comet’s connectivity methods and then export the results for further processing in other environments such as MATLAB. Similarly, the multiverse analysis module operates independently of the specific methods implemented in Comet, meaning it can be flexibly applied to any kind of multiverse analysis pipeline, including those built with user-defined or external functions.

To accommodate different user needs and workflows, Comet provides both a graphical user interface (GUI) and a standard scripting application programming interface (API). While scripting offers greater flexibility and automation, the GUI supports all core functionalities and is particularly useful for exploratory analysis and teaching. After installation, the GUI can be launched directly from the command prompt/terminal (Figure 4).

**Figure 4:**
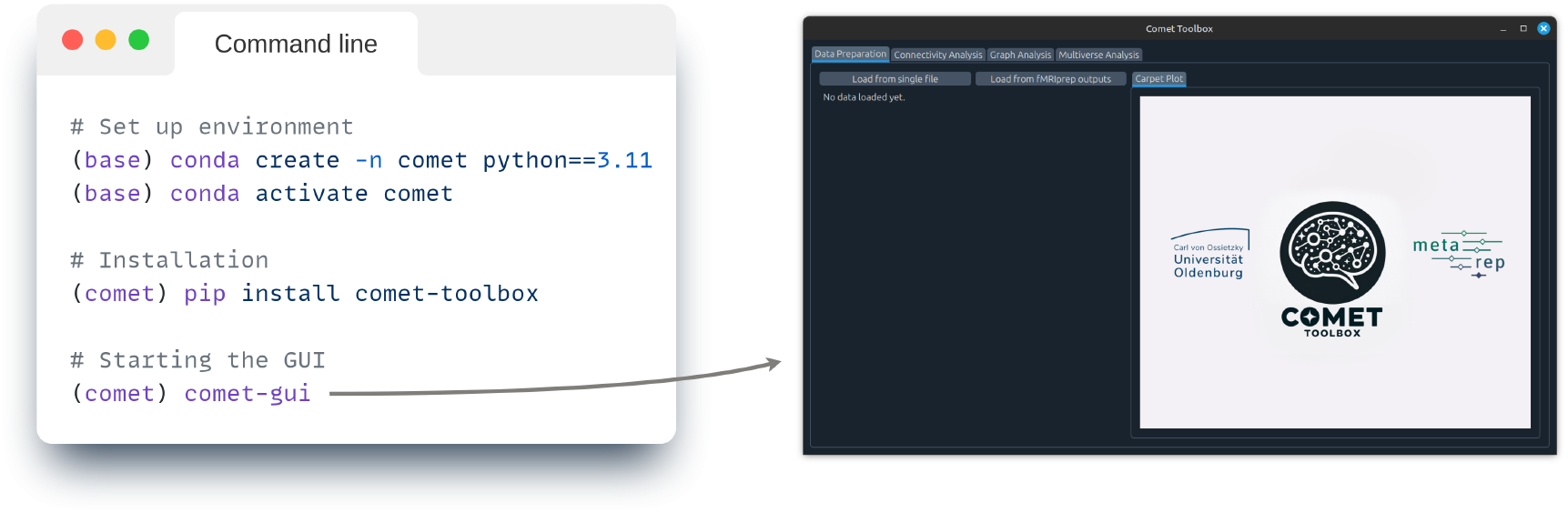
Installation and usage. Comet can be installed via the Python Package Index (PyPI) using pip. Due to the extensive set of dependencies, we strongly recommend installing Comet within a dedicated environment and with Python 3.11 (earlier versions may also be compatible but are not tested). Once installed, the graphical user interface (GUI) can be launched by using the “comet-gui” command.

### 3.1 Graphical User Interface

The graphical user interface currently includes four tabs: Data Preparation, Connectivity Analysis, Graph Analysis, and Multiverse Analysis (as shown in Figure 2).

1. In the *Data Preparation* tab, users can import the desired data. This includes time series data of shape (*N*_time points_ *× N*_brain regions_) in all popular formats (.txt,, .mat, .npy, …), standard fMRI images (.nii, .dtseries), or an entire fMRIprep processed dataset. Depending on the input type, the interface dynamically offers appropriate preprocessing options such as parcellation, temporal filtering, and denoising. The resulting (*N*_time points_ *× N*_brain regions_) time series data can then either be saved to disk or passed forward for connectivity analysis.
2. In the *Connectivity Analysis* tab, users can select any of the included functional connectivity methods, and apply them to the data as prepared in the previous tab. The *Keep in memory* checkbox allows users to keep the calculated dFC estimates in working memory to enable quick switching and comparison between different methods. For computationally expensive dFC methods, a progress bar will be be displayed in the terminal from which the GUI was invoked. Once computed, the connectivity matrices are automatically visualised in the right-hand panel, and users can explore region-to-region connectivity time courses, inspect summary statistics, or export results for external use.
3. In the *Graph Analysis* tab, users can compute a range of network-theoretical measures from connectivity matrices. The GUI accepts input in the form of either static connectivity matrices of shape *N*_brain regions_ *× N*_brain regions_ or dynamic connectivity data with an additional temporal dimension *N*_brain regions_ *× N*_brain regions_ *× N*_time points_. These can be uploaded directly or selected from previously computed results. The interface supports the calculation of commonly used graph metrics such as global efficiency, modularity, degree centrality, clustering coefficient, and community detection.
4. In the *Multiverse Analysis* tab, users can define and evaluate multiverse analyses. This involves specifying a set of decisions, each with multiple options, which are inserted as placeholders into a template script. The template represents the user’s analysis pipeline and can either be written directly in the GUI or exported and edited externally. Once the analysis script is complete and includes all placeholders, it can be reloaded into the GUI for execution and result visualisation. Because this process requires users to write parts of the analysis code themselves, it closely resembles the scripting-based workflow shown in Figure 6.

**Figure 5:**
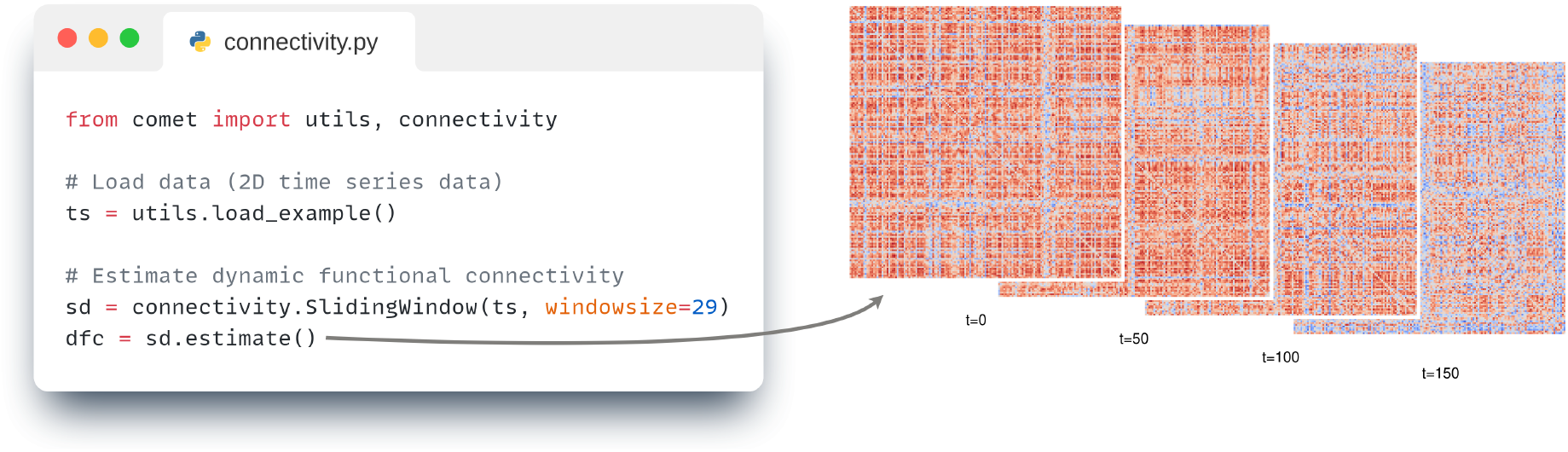
Functional connectivity. Left: Code for estimating (dynamic) functional connectivity. All methods are provided as individual classes, and time-resolved functional connectivity can be estimated through the `.estimate()` method. Right: The resulting sliding window connectivity estimates at four different time points.

**Figure 6:**
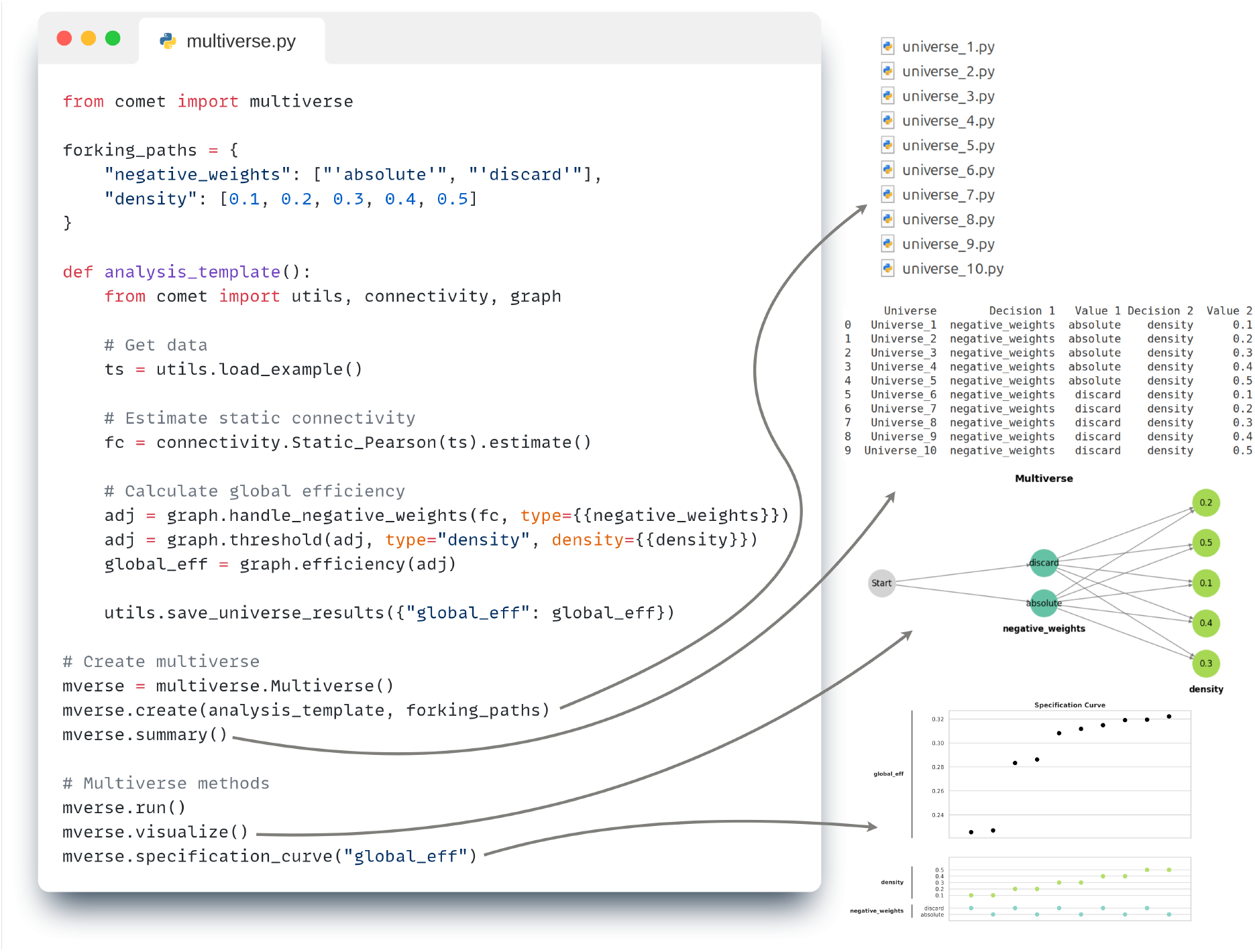
Multiverse analysis workflow in the Comet toolbox. Left: A minimal example of a multiverse analysis script is shown. Forking paths are defined as a dictionary of decision points and their options (e.g., how to handle `negative_weights` and which density thresholds to apply). These are inserted as placeholders (e.g., `{{density}}`) into a template function, which defines the analysis logic (in this case, estimating global efficiency from static functional connectivity). The multiverse is then created using the `.create()` method, and evaluated using the `.run()` method. Right: The toolbox automatically generates Python scripts (e.g., `universe_1.py, universe_2.py`, …) for each valid combination of options, referred to as a *universe*. Each script represents a fully defined analysis pipeline and can be viewed and analysed individually. A table summarises the generated universes, with each row showing the specific options chosen for each decision. The multiverse structure can be visualised as a decision graph, where each branching point represents a forking path. Finally, the results across all universes are summarised using a specification curve, which allow users to identify how different analytic choices affect the outcome (in this case global efficiency).

### 3.2 Scripting API

If more complex analyses are required or a scripting approach is preferred, Comet provides a standard scripting interface that allows direct use of individual modules. For example, as shown in Figure 5, a dynamic functional connectivity method can be used by first importing and instantiating the corresponding class (e.g., `SlidingWindow()`) from the `connectivity` module, and then calling its `estimate()` method. In this example, simulated time series data is loaded from the utils module. A complete list of available classes, methods, and parameters is provided in the online documentation.

Multiverse analyses can be implemented programmatically via the `multiverse` module in standard Python scripts or in a Jupyter Notebook. This approach allows full control over the forking paths, analysis templates, and optional configurations. Forking paths (i.e., decision points with multiple possible options) are defined in a Python dictionary. Each combination of options corresponds to a unique analysis pipeline, referred to as a *universe*.

An example using two forking paths is shown in Figure 6. The first forking path (`negative_weights`) defines two options for handling negative values in a connectivity matrix. The second path (`density`) specifies five possible thresholds for graph construction. These options are represented in the analysis script by placeholders written in double curly brackets, such as {{`density`}}. During multiverse generation, these placeholders are automatically replaced by each of the defined options, allowing the toolbox to generate a complete analysis script for all valid combinations of decisions.

Once the template and forking paths are defined (in this simple example a pipeline for estimating global efficiency from static functional connectivity), the multiverse is created using the `.create()` method, which generates individual Python scripts for each universe (e.g., `universe_1.py, universe 2.py`, etc.). These scripts can be executed directly using the `.run()` method, either sequentially or in parallel. The multiverse can be created, analysed, and visualised, with the most important built-in functions being:

`multiverse.Multiverse()` Initializes the multiverse object,

`.create()` Generates universes and corresponding scripts,

`.summary()` Provides an overview of decisions and universes,

`.visualize()` Plots the structure of the multiverse,

`.run()` Executes the universes (in full or in part),

`.specification_curve()` Visualises variability across universes,

`.get_results()` Get all (or parts) of the results,

In the example shown, the two forking paths with 2 *×* 5 options yield ten unique universes. Each universe script contains a fully specified analysis pipeline corresponding to one combination of decisions. The multiverse structure is shown as a branching graph, and the results are summarised in a specification curve. More advanced features, such as defining invalid decision combinations or specifying decision orders, are supported via an optional configuration dictionary. Detailed tutorials are available in the documentation.

## 4 Analysis Example

We showcase the toolbox and multiverse analysis workflow using simulated fMRI data. For this, the SimTB package (Erhardt et al., 2012) was used to generate example time series data from 10 nodes whose connectivity structure alternates between two discrete states in a block-like design. The simulation consists of 100 blocks of 7.2 seconds each, with one of the two states randomly selected as the dominant connectivity pattern per block. Each node independently has a 20% chance of experiencing a unique event at any time point and is further affected by Gaussian noise, adding to the complexity of the signal and creating a more challenging scenario for the analysis. The Jupyter Notebook for this example (as well as many other examples) can be found in the tutorial scripts for the toolbox, and we encourage interested readers to explore this resource.

The aim of this example is to demonstrate how the toolbox handles multiverse analyses in the domain of dFC analyses. For this, we constructed a multiverse analysis containing 240 (5 *\** 2 *\** 2 *\** 2 *\** 3 *\** 2) universes which include analytical decisions for dFC estimation, graph network construction, and the prediction model. A summary of these forking paths is provided in Table 2. We emphasize that this example is intended as a practical illustration of the toolbox functionality and we do not aim to interpret the outcomes of these decisions.

**Table 2:** Decisions and options for the example analysis.^a^

| Decision | Option |
| --- | --- |
| dFC method | Tapered Sliding Window Correlation<br>Flexible Least Squares<br>Multiplication of Temporal Derivatives<br>Phase Synchronization<br>Spatial Distance |
| Temporal lag | 6 TR<br>10 TR |
| Graph density | Strongest 50%<br>Strongest 25% |
| Graph weights | Weighted<br>Binarized |
| Graph measure | Local Efficiency<br>Clustering Coefficient<br>Participation Coefficient |
| SVC kernel | Linear<br>Radial Basis Function |

<sup>a</sup> The multiverse consists of 240 unique combinations of decisions ( $5 \times 2 \times 2 \times 2 \times 3 \times 2$ ), referred to as universes. For each universe, a support vector classifier (SVC) was trained to predict the dominant connectivity state of each simulated block, based on features extracted from the corresponding time window. Results of these classification analyses are summarised in Figure 7.

In detail, we applied six distinct dFC methods (Sliding Window Correlation, Flexible Least Squares, Multiplication of Temporal Derivatives, Phase Synchronization, Spatial Distance), each grounded in different theoretical assumptions. The simulated time series consisted of alternating blocks, each corresponding to a specific ground truth connectivity pattern. The TR (repetition time) was set to 0.72 seconds, consistent with typical modern multiband acquisition. To align with this design, we divided the resulting dFC time series into 100 blocks. Each block was treated as a classification instance, corresponding to one simulated connectivity state. To account for the temporal delay introduced by the hemodynamic response in BOLD fMRI, we introduced a temporal shift of either 6 or 10 TRs to the block onsets before extracting the features.

For each block, we averaged the dFC estimates across its time points to obtain a single connectivity matrix. Graph networks were then constructed from these matrices based on two binary decisions: (1) whether to retain the top 50% or top 25% of edges (graph density thresholding), and (2) whether to binarise the resulting graph or retain edge weights. From each graph, we computed three node-level graph-theoretic measures: the clustering coefficient, participation coefficient, and local efficiency. These measures served as features for classification.

For the prediction model, we trained a support vector classifier (SVC) to predict the connectivity state of each block, comparing two kernel types: linear and non-linear (radial basis function). Classification performance was assessed using 5-fold cross-validation. The results were visualised using a specification curve (Figure 7), which arranges all pipeline configurations by their classification accuracy in ascending order. This provides a comprehensive view of how analytic choices affect model performance.

**Figure 7:**
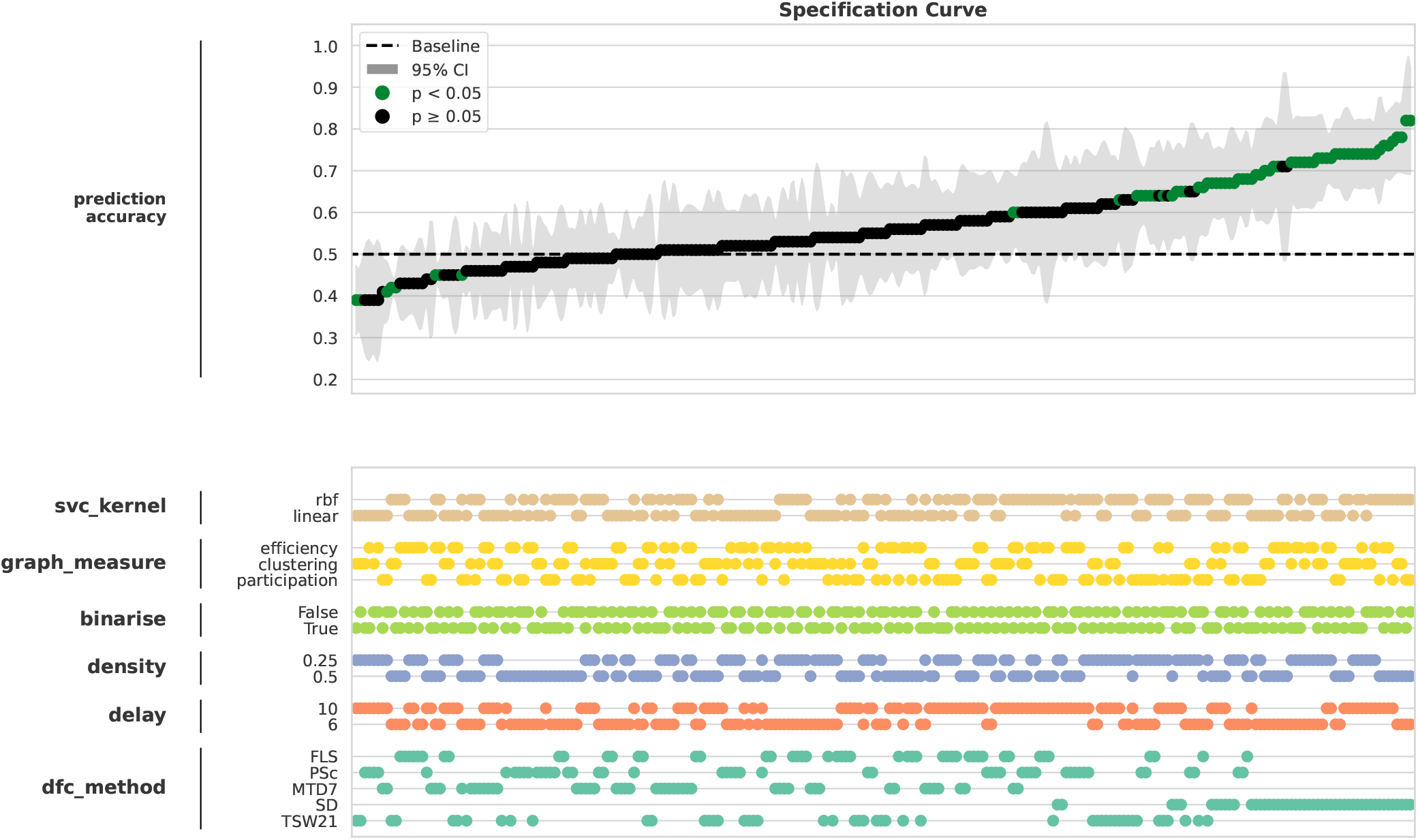
Specification curve for the example analysis. The classification accuracies for the individual universes are displayed in ascending order. The upper plot displays the classification accuracies with their 95% confidence intervals resulting from 5-fold cross validation. The green dots indicate accuracies significantly above chance (uncorrected). The bottom plot displays the individual decisions chosen in the analysis pipeline leading to the corresponding classification accuracy above.

Notably, since our synthetic data included two clearly separable ground-truth states, above-chance classification was expected. Yet, the specification curve shows that many analysis pipelines yielded accuracies near chance, with overlapping 95% confidence intervals. Only one dFC method (Spatial Distance; SD) consistently enabled accurate classification. This highlights how analytic choices might obscure real effects, leading to false negatives. Multiverse analysis thus serves not only as an aggregative, but also as an exploratory framework, revealing systematic patterns in the decision space that can inform future methodological choices and guide validation on independent datasets.

## 5 Discussion

The present work introduced the Comet toolbox, a dedicated framework for conducting multiverse analyses within network neuroscience. We believe the toolbox addresses a critical need in a rapidly growing research area, where multiverse analysis is increasingly used to assess the robustness of findings in fMRI studies. For example, Luppi et al. (2024) conducted a large-scale multiverse analysis of data-processing pipelines to promote more consistent practices in functional connectomics, while Demidenko et al. (2024) applied multiverse analysis to explore the influence of preprocessing decisions in the context of the monetary incentive delay task. In parallel, more conceptual contributions such as Lefort-Besnard et al. (2025) have framed multiverse analysis as a form of same-data meta-analysis and proposed integration strategies for evaluating analytic variability, although these efforts have so far been limited to test statistic maps from activation studies.

At the same time, broader methodological needs in network neuroscience remain unresolved. For example, Fekonja et al. (2025) recently outlined nine key roadblocks that hinder clinical adoption of network-based methods, including difficulties in constructing accurate and interpretable networks, the computational complexity of dynamic connectivity analysis, and the lack of accessible, user-friendly tools suitable for clinical workflows. Comet aims to address several of these barriers by providing (i) a systematic framework for exploring analytic flexibility, (ii) integrated modules for dynamic functional connectivity and graph-theoretic analysis, and (iii) a dual interface (graphical and script-based) that supports both technical and clinical users.

### 5.1 Comparison to existing software

Comet aims to address a specific methodological gap in network neuroscience by combining dynamic functional connectivity (dFC) estimation, graph-theoretical analysis, and multiverse workflows within a single, unified platform. While various toolboxes are available for individual aspects of this workflow, few provide an integrated framework that supports both methodological diversity and systematic multiverse analysis. While not exhaustive, the following subsections outline related toolboxes and approaches to situate Comet within the current software landscape.

Many existing packages implement one or a few specific dFC methods, often developed alongside individual publications. These include, for example, the `timecorr` package for kernel-based correlation (Owen et al., 2021), the Dynamic Correlation toolbox for GARCH-based models (Lindquist et al., 2014), or implementations of methods such as Leading Eigenvector Dynamics (Cabral et al., 2017; Olsen et al., 2022). While these tools provide valuable methodological contributions, their individual scope, and in some cases their tight integration with publication specific workflows, can limit their reusability and accessibility.

Some Python and MATLAB-based toolboxes offer broader functionality for dFC analysis. For example, the Python toolboxes `teneto` (Thompson et al., 2017) and `pyDFC` (Torabi et al., 2024) support multiple dFC methods and allow for modular analysis pipelines. However, neither is focused on complex multiverse workflows, and both rely on internal data formats that may require additional steps to integrate with widely used neuroimaging toolchains such as `nilearn`. In MATLAB, the DynamicBC toolbox (Liao et al., 2014) and the CONN toolbox (Nieto-Castanon, 2020) include support for dFC estimation, but do not provide explicit tools for multiverse analysis or analytic comparison across parameter spaces. In contrast, Comet is designed to support structured exploration of analytic variation and integrates both various approaches of dFC estimation and graph-based analyses in a way that is directly compatible with multiverse workflows. Notably, Alteriis et al. (2025) proposed the Dysco framework as an alternative strategy for reducing analytic uncertainty by combining multiple dFC methods into a single integrated approach. While this differs from the multiverse framework used in Comet, it reflects a shared aim of improving robustness by addressing uncertainty across methodological choices.

Other packages focus on full pipeline integration, from preprocessing to graph-theoretic analysis. Examples include GRETNA (Wang et al., 2015), which supports static and dynamic connectivity analysis with graph metrics, and ACTION (Fang et al., 2025), which provides a GUI-based pipeline for static functional connectivity and machine learning models. While these toolboxes offer end-to-end functionality, they do not incorporate multiverse analysis features, and typically support only a limited selection of connectivity and graph-theoretic methods. Comet differs in its explicit support for user-defined decision trees, flexible graph construction options, and scripting interfaces for multiverse evaluation.

Finally, outside the domain of dFC and network neuroscience, a number of general-purpose multiverse analysis tools have emerged. These include Boba, multiverse, and specr R packages (Y. Liu et al., 2020; Masur & Scharkow, 2020; Sarma et al., 2023), which are designed for behavioural and statistical data in Python and R, respectively. Other packages focus on specific experimental domains, such as fear conditioning (Lonsdorf et al., 2022) or the visualisation of multiverse results (Sarma et al., 2024). While these tools offer important contributions to the broader discussion on analytical flexibility, they are typically not tailored to the complexity of neuroimaging workflows. In contrast, Comet was developed specifically to address the needs of fMRI and network neuroscience research, where analysis pipelines often span multiple stages of data processing and modelling decisions.

In summary, while many existing toolboxes offer powerful features for specific parts of the analysis pipeline, none provide the methodological diversity, extensibility, and integration needed to support multiverse analyses in the context of dFC and graph-theoretical neuroscience. Comet is designed to bridge this gap by combining established and novel methods for dFC, flexible graph analysis tools, and a dedicated multiverse engine, all within a single, accessible framework.

### 5.2 Future Directions

Comet is an actively developed toolbox and will continue to evolve over time. At present, it provides a flexible framework for multiverse analysis and supports a wide range of dynamic functional connectivity and graph-theoretic methods. Looking ahead, several possible directions for expansion may further enhance the utility and scope of the toolbox. Beyond its current primary use in assessing the robustness of results across analytic choices (e.g., via specification curve analysis; Simonsohn et al. (2020)), future developments could explore additional conceptual perspectives. For instance, multiverse analysis has recently been proposed as a tool for identifying covariates that substantially influence outcomes (Bowring et al., 2022), or as a framework for supporting abductive reasoning and inferential decision-making in complex analyses (Engzell & Mood, 2023). Related efforts in the fMRI domain are beginning to investigate approaches for statistical inference and multiverse integration (Girardi et al., 2024; Lefort-Besnard et al., 2025; Ozenne et al., 2025), and could inform future methodological modules in Comet. Techniques such as active learning have also been suggested to optimise multiverse exploration when a full combinatorial search is infeasible (Dafflon et al., 2022). While these developments remain at the research frontier, they highlight promising avenues for extending Comet’s functionality. Future versions of the toolbox might also incorporate temporal network measures that go beyond static snapshot topology, as e.g. described by Thompson et al. (2017). Additionally, tools for studying inter-individual differences from a psychometric perspective are not yet integrated, despite their relevance for personalised network neuroscience (Forkel et al., 2022). Overall, we see Comet not as a static software package, but as a flexible and extensible platform that can adapt to these evolving needs of the neuroimaging community.

### 5.3 Conclusion

The Comet toolbox provides a unified yet flexible platform for dynamic functional connectivity (dFC) and network analysis, enabling researchers to systematically explore how methodological choices influence study outcomes. It integrates a broad range of dFC methods alongside graph-theoretical tools. Multi-verse analysis addresses the growing concerns about robustness and reproducibility in neuroimaging by facilitating structured comparisons across diverse analytical pipelines. The toolbox outputs are designed to support a deeper understanding of how analytic variability affects research findings.

Comet is accessible through both a graphical user interface and a simple scripting API, making it suitable for users with varying levels of programming expertise. By supporting both conventional and multiverse-style workflows, the toolbox serves as a valuable resource for enhancing transparency, flexibility, and reproducibility in neuroimaging research. The toolbox and documentation are available at: https://github.com/mibur1/comet.

## Ethics

All data used in this study were entirely synthetically generated.

## Data and Code Availability

The data, code, and documentation for the toolbox are publicly available at https://github.com/mibur1/comet. The documentation website provides comprehensive guidance for installation and usage, offering a broad range of tutorials and API information.

## Author Contributions

**Micha Burkhardt:** Conceptualization, Data curation, Formal Analysis, Investigation, Methodology, Software, Visualization, Validation, Writing - original draft, Writing - review and editing. **Carsten Gießing:** Conceptualization, Funding acquisition, Project Administration, Supervision, Writing - review and editing

## Acknowledgements

This work was supported by a grant from the German Research Foundation (DFG) to Carsten Gießing (GI 682/5-1 and GI 682/5-2), Christiane Thiel (TH 766/9-1 and TH 766/9-2) and Andrea Hildebrandt (HI 1780/7-1 and HI 1780/7-2) as part of the DFG priority program “META-REP: A Meta-scientific Programme to Analyse and Optimise Replicability in the Behavioural, Social, and Cognitive Sciences” (SPP 2317).

## Supplementary Material

The Comet toolbox includes a large array of dynamic functional connectivity methods, which we briefly describe in the following subsections.

### Continuous Connectivity Methods

#### Sliding Window Correlation

To date, sliding window correlation (SWC) is the predominant method for assessing dynamic functional connectivity (Lurie et al., 2020; Shakil et al., 2016). The SWC method works by sliding a window with a specified size, shape, and step size over the time series and calculatin the pairwise Pearson correlation between the resulting subsets of time series. Given two time series *x*(*t*) and *y*(*t*), the SWC for a window *i* corresponds to the Pearson correlation coefficient within that window:

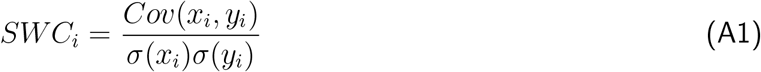

Here, *x*_*i*_ and *y*_*i*_ are the subset of *x*(*t*) and *y*(*t*) within the *i*^*th*^ window, *Cov*(*x*_*i*_, *y*_*i*_) is the covariance between *x*_*i*_ and *y*_*i*_, and *σ* denotes the standard deviations of *x*_*i*_ and *y*_*i*_, respectively. Considering data for *N* brain regions (*N*_*nodes*_), this creates *N*_*nodes*_ *×* (*N*_*nodes*_ − 1)*/*2 = *N*_*edges*_ unique dynamic functional connectivity (dFC) estimates for each window, which are usually summarized in symmetric connectivity matrices of shape *N*_*nodes*_ *×N*_*nodes*_. Inspecting the temporal fluctuations of values in these matrices reveal the time course of dFC estimates, which, due to the low-pass properties of windows, are smoothed time courses reflecting the time-varying connectivity between each pair of brain areas.

Sliding window correlations are a mathematically and computationally simple, but also come with significant caveats. The most discussed issue stems from the choice of the window length, which controls the a trade-off between temporal sensitivity and accuracy. Small window sizes increase the risk of spurious fluctuations in dFC due to noise and also hinder a reliable estimation of the Pearson correlation, while large windows result in the loss of temporal variations in the signal.

#### Multiplication of Temporal Derivatives

Multiplication of Temporal Derivatives (MTD) was first introduced by Shine et al. (2015) as a method to overcome the aforementioned shortcomings of the sliding window. The method first calculates the temporal derivative *dt* for each time series *x*(*t*) at each time point *t*:

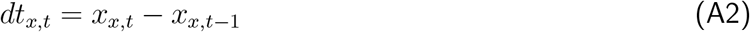

Then, the coupling between each pair of derivative time series *dt*_*x*_ and *dt*_*y*_ is computed by multiplying them at each time point *t* and dividing by the product of their standard deviation.

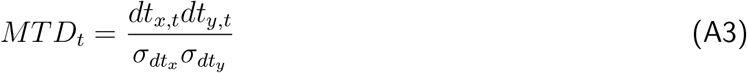

Thus, MTD provides a measure of functional coupling between time series. If signals increase or decrease together MTD will be positive, if signals change in opposite direction MTD will be negative. As a frame-wise estimate operating on individual time points, MTD can, however, be susceptible to high-frequency noise. Thus, Shine et al. (2015) suggest to use temporal smoothing by using a simple moving average for the MTD time series with a window size of 7 time points.

#### Jackknife Correlation

The Jackknife Correlation (JC) method (Richter et al., 2015) computes the covariance of two time series *x*(*t*) and *y*(*t*) at time point *t* by using all time points except *t*. This therefore creates estimates of covariance on a single trial basis and can intuitively be seen as an inverse sliding window of length one:

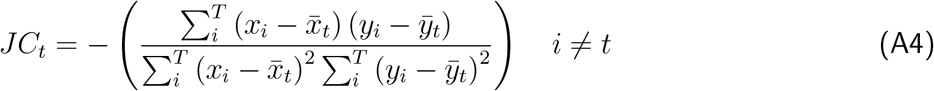

The Jackknife Correlation *JC*_*t*_ is calculated for each time point *t*. The leave-1-out process results in an inversion of the direction of connectivity estimates, which is corrected by the minus sign. The primary advantage of the JC method is that it addresses the aforementioned challenges associated with small window sizes as mentioned above. In this context, the method has previously been described as an optimal sliding window, as it avoids the trade-off between temporal sensitivity and accuracy since the calculation of JC at each time point *t* is based on *N*_*time*_ − 1 data points, with *N*_*time*_ being the length of the time series (Thompson et al., 2018).

It is to note that this method is also not without its limitations. First, JC is highly sensitive to high-frequency noise in the data. This could, however, be reduced by excluding more than one time point for each estimate, resulting in a leave-n-out procedure. Second, it only provides relative estimates of connectivity that fluctuate around the static connectivity of each subject. And third, it leads to a compression of variance proportional to the length of the time series as each estimate differs by just one data point. To that avail, Thompson et al. (2018) propose further processing steps like standardisation of the resulting JC time series for direct comparison between subjects.

#### Edge-centric Connectivity

Edge-based functional connectivity was recently proposed by Faskowitz et al. (2020) and more closely described in Betzel et al. (2023) as a framework for functional connectivity with a focus on edges instead of nodes. Briefly, the framework introduces edge time series (eTS) and edge functional connectivity (eFC).

Given two z-scored time series *x*(*t*) and *y*(*t*), the eTS is defined as the element wise product of the original time series:

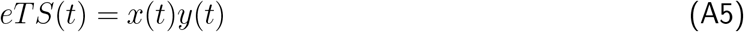

This process is done for all pairs of brain regions, resulting in an eTS of shape *N*_*edges*_ *× N*_*time*_. As the edge dimension is identical to the unfolded triangle of a standard connectivity matrix, this measure can be seen as an unnormalised and unsmoothed version of the Multiplication of Temporal Derivatives method and is also highly similar to the Jackknife Correlation Method. Estimating eTS thus requires no parameters and provides FC estimates at the resolution of individual time points.

A notable feature of eTS is that their normalized sum across time yields the Pearson correlation coefficient:

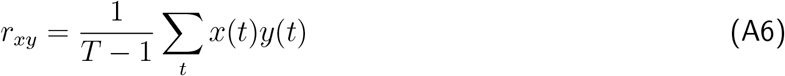

which means that they can be interpreted as an exact decomposition of static functional connectivity. Based on the eTS, eFC can then be derived by quantifying the similarity between pairs of edge time series. For two eTS, *r*_*xy*_(*t*) and *r*_*uv*_(*t*), the eFC is defined as:

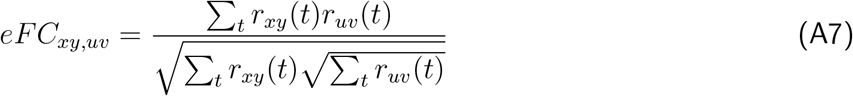

This results in a single edge connectivity matrix of shape (*N*_*edges*_*×N*_*edges*_), which can be memory intensive for larger parcellations.

#### Spatial Distance

The previously introduced methods all use temporally adjacent time points to estimate the covariance between brain areas. Spatial Distance (SD) (Thompson et al., 2017) differs from that approach by using time points that have a similar spatial profile. Initially, a weight vector is computed for each time point *t* by calculating the spatial distance (e.g., Euclidean distance) to all other time points. As introduced by Thompson et al. (2017), Comet implements the inverse of the Euclidean Distance (*ED*) between the BOLD signal amplitudes at time point *t* and all other time points *u*:

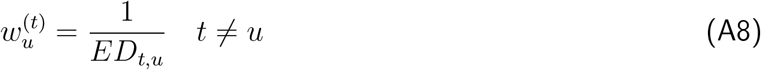

Assuming a time series containing 100 time points, SD will then create a vector of length 100 for each time point, with each vector containing the spatial distance of the activity pattern from the corresponding time point to all other time points. Each weight vector is then normalized, scaling the weights between 0 and 1, with the self-weight at time point *u* being set to 1. Subsequently, the functional connectivity between two brain regions at time point *t* can be calculated as the weighted Pearson correlation coefficient, utilizing the weight vector 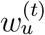. SD has shown to be effective in capturing dynamic changes in connectivity by prioritizing spatial similarities. However, it may be influenced by the choice of distance metric and normalization process (Thompson et al., 2017).

#### Flexible Least Squares

Flexible Least Squares (FLS) is a time-varying flexible regression approach introduced by Kalaba and Tesfatsion (1989), which gained popularity in the domain of dFC through its inclusion in the DynamicBC toolbox (Liao et al., 2014). FLS estimates changing relationships between time series through a *β* coefficient that varies over time to minimize data fitting errors. It is a data-driven and distribution-free method that accounts for both measurement and dynamic errors to capture evolving associations between the time series. First, a general regression model is assumed in the form of:

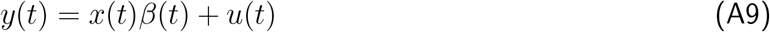

In this equation, *x*(*t*) and *y*(*t*) are two time series, *β*(*t*) determines the strength of the covariance between them, and *u*(*t*) is the approximation error. Opposed to static linear regression, which estimates a single beta coefficient, FLS estimates a time-varying sequence of betas by minimizing 1) a measurement error between the actual and estimated observation at each time point, and 2) a dynamic error which represents the discrepancy due to incorrect specification of the state transition equations. The measurement fit error is described by:

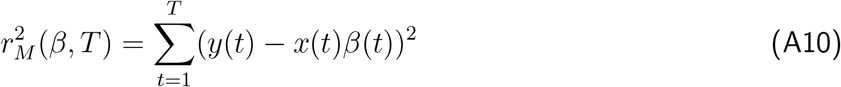

where *x*(*t*) and *y*(*t*) represent two time series of length *T* at time *t* and *β* denotes the model fit coefficient vector. The dynamic error is described by:

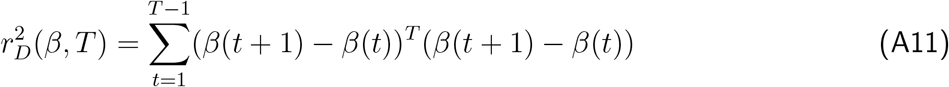

This dynamic error also has the objective of smoothing the *β*(*t*) coefficients by penalizing large variations between time points and thus preventing overfitting to noise. Both errors are then combined into a single cost-incompatibility function to estimate the time-varying *β*(*t*) sequence:

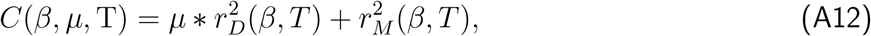

This function is then minimized through ordinary least squares, with *µ* forming a weighting parameter between the measurement fit error 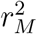 and the dynamic error 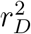. The resulting *β* coefficient between two signals at time point *t* is then interpreted as the functional connectivity. As previously mentioned for JC and SD, the FLS method also has the advantage of not relying on a specified window length and thus avoids the aforementioned issues that come with this choice. However, as the optimization procedure has to be performed for each pair of signals, the method is computationally demanding.

#### Dynamic Conditional Correlation

Dynamic Conditional Correlation (DCC) (Engle, 2002) is an extension of GARCH models predominantly used in economics and finance. DCC models time-varying correlation between time series by adapting to transient fluctuations in data. Its use was recently introduced in the context of fMRI analyses by Lindquist et al. (2014), who found DCC to outperform previously established methods for estimating dFC. A GARCH model is described through the following equations:

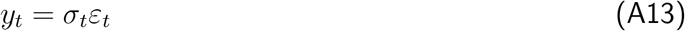

Here, it is first assumed that a time series *y*_*t*_ can be expressed as the product of a time-varying variance term *σ*_*t*_ and random variable *ε*_*t*_ *∼* 𝒩 (0, 1) referred to as innovation. In a GARCH(1,1) model, which takes into account a single autoregressive term and a single moving average term, the conditional variance is then expressed as:

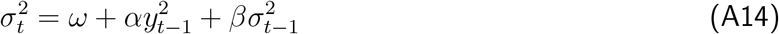

with parameters *ω >* 0, *α, β ≥* 0 and *α, β <* 1. *α* modulates the influence of the squared previous observation 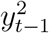 and *β* modulates the influence of the previous estimate of the conditional variance 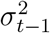. These parameters are then optimized through maximum likelihood estimation to best match the time series.

It should be noted that this can be generalized to GARCH(p,q) processes, with *p* denoting the number of previous observations and *n* denoting the number of previous estimates of the conditional variance to be included in the model. This could especially prove fruitful when using data acquired with a small TR, however this has yet to be explored in the context of fMRI dFC analyses.

As an extension to GARCH, the DCC method has been shown to be superior to other multivariate GARCH models that try to estimate dynamic correlations (Engle, 2002). To show the general working of the model, *y*_*t*_ = *ε*_*t*_ is considered as a bivariate time series with mean zero and a conditional covariance matrix Σ_*t*_. DCC is then expressed through the following set of equations:

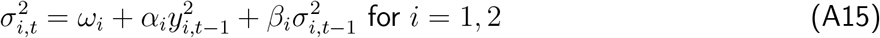

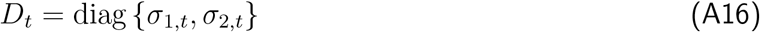

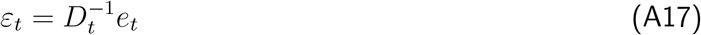

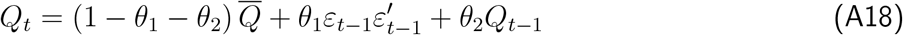

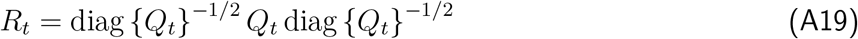

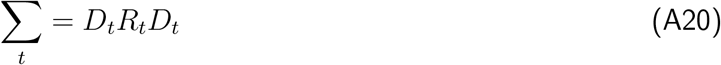

These equations describe two general steps. First, univariate GARCH(1,1) models are fit to each time series in *y*_*t*_ (Eq. 12), and standardized residuals are calculated (Eq. 14). Second, *Q*_*t*_, a non-normalized version of the time-varying correlation is derived using an EWMA method on standardized residuals (Eq. 15). The matrix 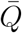 is the unconditional covariance of *ϵ*_*t*_, with scalars *θ*_1_, *θ*_2_ constrained as 0 *< θ*_1_+*θ*_2_ *<* 1. They are adjusted for scale to form a valid correlation matrix (Eq. 16), and the time-varying covariance matrix is established (Eq. 17). The model parameters, including *ω*_1_, *α*_1_, *β*_1_, *ω*_2_, *α*_2_, *β*_2_, *θ*_1_, *θ*_2_, are estimated through a bifurcated process: initially, by estimating time-varying variances for each time series, and subsequently by utilizing standardized residuals to estimate dynamic correlations as shown by (Engle, 2002).

The strength of the DCC model is its capability to detect rapid changes in data correlation structures with relatively few assumptions, and it has been shown to outperform sliding window techniques but also faces challenges such as computational complexity, potential numerical issues during optimization, as well as a risk of overfitting to noise (Lindquist et al., 2014). The latter is particularly problematic in fMRI data, where the signal-to-noise ratio is generally low.

#### Phase Synchronization

Phase Synchronization (PS) models originate from the field of physics, and have initially been used to examine the behaviour of weakly coupled oscillators (Rosenblum et al., 1996). The general idea behind PS methods is to convert a real signal into its complex analytic version to obtain instantaneous phase and amplitude information, and then derive pairwise coherence estimates. While this can be achieved in various ways, phase synchronization methods usually apply the Hilbert transform, which is described as follows:

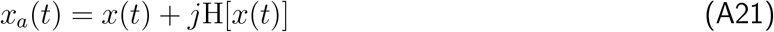

Here, the real signal *x*(*t*) is translated into its analytic form *x*_*a*_(*t*) with *H* being the Hilbert transform and *j* being the imaginary unit. The instantaneous phase is then defined as:

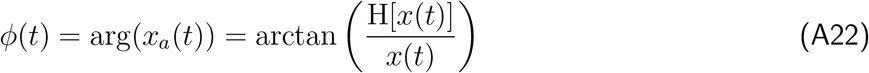

Given the instantaneous phase *ϕ*_*i*_ and *ϕ*_*j*_ of two analytic signals, their coherence can be estimated in multiple ways. The toolbox includes two different approaches, namely the PSc (Honari et al., 2021; Maltbie et al., 2022) and and PSp (Honari et al., 2021). These approaches are implemented as follows:

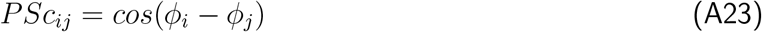

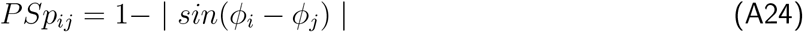

Here, instantaneous PS is estimated by either taking the cosine of the instantaneous phase difference between a pair of signals, or by subtracting the absolute of the sine of the instantaneous phase difference from one. This results in different measures of PS, as it produces a coherence matrix with estimates ranging from -1 (anti-phase) to 1 (in phase) for PSc, and coherence estimates ranging between 0 (no phase coherence) and 1 (perfect phase coherence) for PSp.

The main limitation of these phase synchronization methods is that the Hilbert transform only produces meaningful results on a narrowband signal, and the fMRI time series therefore needs to be filtered accordingly. While fMRI studies usually filter signal in the range of 0.01-0.1 Hz, this is band is still too broad and literature suggests smaller bands (e.g., 0.03-0.07 Hz) are required to obtain meaningful results (Glerean et al., 2012; Honari et al., 2021; Sheng et al., 2023).

#### Leading Eigenvector Dynamics

Leading Eigenvector Dynamics (LeiDA) extends the idea of the PS method by deriving the leading eigenvector from the coherence matrix and is implemented as described by Cabral et al. (2017) and Olsen et al. (2022). As previously described for the PS method, LeiDA relies on the Hilbert transform to extract the instantaneous of an analytic signal. From the instantaneous phase, a coherence matrix *C* is then computed analogous to PSc with *i* and *j* indexing over brain regions:

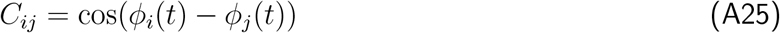

The eigendecomposition of this coherence matrix then provides its eigenvectors and eigenvalues. The leading eigenvector, which corresponds to the matrix’s largest eigenvalue, is of particular interest:

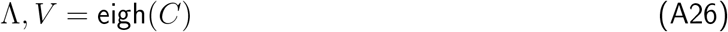

where Λ is the diagonal matrix of eigenvalues and *V* are the corresponding eigenvectors with the leading eigenvector *V*_1_. *V*_1_ is then interpreted as a measure of functional connectivity, with nodes of equal sign being considered as coherent, and the size of *V*_1_ representing the strength of this coherence. This reduces the dimensionality of each dFC estimate from *N*_*edges*_ to *N*_*nodes*_, which is usually the preferred output for the LeiDA method. However, a full connectivity matrix can be reconstructed at each time point *t* as the outer product of the leading eigenvectors with their transpose, which has been shown to still explain most of the variance of the original coherence matrix (Cabral et al., 2017):

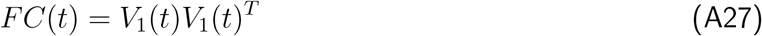

#### Wavelet Coherence

Wavelet Coherence (WCOH) is implemented as described by Billings et al. (2021). In contrast to phase synchronization methods which focus on a a single, narrow frequency band, WCOH makes use of the continuous wavelet transform to transform the real signal *f* (*t*) into time-frequency spectrograms and calculates pairwise coherence between them. The wavelet transform is given by:

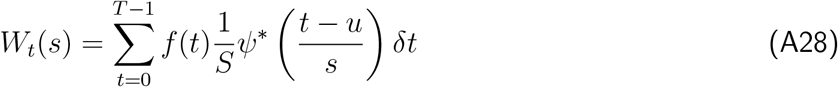

Here, *W*_*t*_(*s*) represents the wavelet coefficient at time *t* and frequency scale *s*. The function *ψ*^*\**^ is the complex conjugate of the wavelet function, while *u* serves as the translation parameter that facilitates the shifting of the wavelet across the time series. The squared wavelet coherence 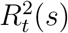 between two time series *X* and *Y* can then be computed as:

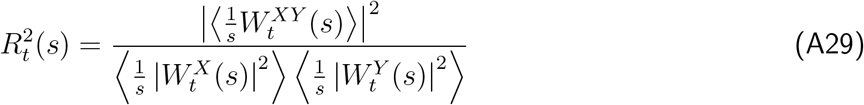

In this equation, 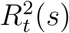 denotes the squared coherence at time *t* and scale *s*. The term 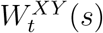 is the cross-wavelet transform between *X* and *Y*, while 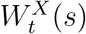 and 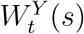 are their individual wavelet transforms. The operator ⟨·⟩ indicates a smoothing operation in both time and scale, and the factor *s*^−1^ serves as a normalization in the wavelet transform.

While it is in theory possible to perform subsequent analyses for separate frequency bands, it is often useful to obtain a unified dFC measure. To achieve this, Billings et al. (2021) proposed to take a weighted average over all scales:

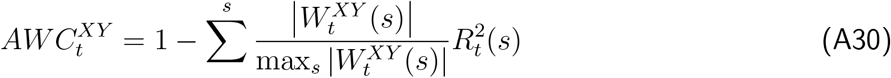

Here, the cross-wavelet power 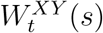 is normalized and used for the calculation of a weighted mean of 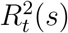 over all frequency bands, resulting in an average wavelet coherence (*AWC*) time series for each pair of signals.

In contrast to the previously introduced methods of PS and LeiDA, WCOH has the benefit of being applicable beyond narrow-band signals. However, a peculiarity with WCOH are ‘cone of influence artifacts’ which refer to the distortions at the edges the spectrograms, caused by the finite length of the data. A common approach to address this is by removing the outer scales and time points, but this leads to a loss of potentially important data, especially in the case of the latter. Moreover, WCOH necessitates variable smoothing across both time and frequency domains (Grinsted et al., 2004), making it the most heavily smoothed method in the toolbox. This needs to be considered when fast fluctuations in dFC are of interest.

### State-based Connectivity Methods

#### Sliding Window Clustering

The Sliding Windlow Clustering (SWC) approach is included as described by Allen et al. (2014) and as implemented by Torabi et al. (2024). Briefly, this approach begins by constructing dynamic functional connectivity (dFC) matrices using a sliding-window technique. Each time point from the dFC matrices is vectorized into a feature vector representing pairwise ROI relationships. A two-level k-means clustering is then applied: initially clustering the individual feature vectors (subject-level), and subsequently clustering the resulting centroids to define FC states. The centroids are reshaped back into their original ROI × ROI format to form the FC matrices for each state. Hyperparameters of this method include the sliding-window configuration, the number of subject-level clusters, and the number of FC states.

#### Coactivation Patterns

The Coactivation Patterns (CAP) approach is included as described by X. Liu and Duyn (2013) and as implemented by Torabi et al. (2024). The CAP method is a point-process analysis that focuses on individual time points with minimal assumptions. The original implementation uses activation thresholding within a seed region to extract significant time points and averages the corresponding BOLD signals. An extended version employs k-means clustering directly on the BOLD time series to derive distinct FC states. The method typically involves a two-stage clustering process, first clustering individual time points at the subject level, and second clustering the resulting centroids at the group level. The state FC matrices are obtained by computing the outer product of the centroids.

#### Continuous Hidden Markov Model

The Continuous Hidden Markov Model (CHMM) approach is included as described by Vidaurre et al. (2017) and as implemented by Torabi et al. (2024). The CHMM method models the continuous BOLD time series as observation sequences in a hidden Markov framework, using a Gaussian observation model. In this approach, the BOLD data are directly used to infer a sequence of hidden states, each represented by a multivariate Gaussian distribution. The state FC matrices are derived from the covariance matrices of these Gaussian models.

#### Discrete Hidden Markov Model

The Discrete Hidden Markov Model (DHMM) approach is included as described by Ou et al. (2015) and as implemented by Torabi et al. (2024). In the DHMM method, a discrete hidden Markov model with a categorical observation model is applied to an initial clustering-based analysis. The clustering yields discrete state sequences which serve as the observation input for the DHMM. The model then infers hidden states corresponding to FC states, with state FC matrices obtained by averaging the connectivity matrices assigned to each state from the clustering output.

#### Window-less

The Window-less (k-SVD) approach is included as described by Yaesoubi et al. (2018) and as implemented by Torabi et al. (2024). This method estimates dominant linear patterns in the BOLD time series using a sparse dictionary learning algorithm, specifically the k-SVD algorithm. Each dictionary element represents a dominant linear pattern corresponding to an FC state. A hard sparsity constraint is applied so that each time point is represented by a single dominant dictionary element. The state FC matrices are computed as the outer products of these dictionary elements.

## Footnotes

1 https://docs.python.org/3/library/venv.html

2 https://conda.io/projects/conda/en/latest/index.html

